# The conspicuousness of the toxic *Heliconius* butterflies across time and habitat

**DOI:** 10.1101/662155

**Authors:** Denise Dalbosco Dell’Aglio, Jolyon Troscianko, Martin Stevens, W Owen McMillan, Chris D Jiggins

**Affiliations:** Butterfly Genetics Group, Department of Zoology, University of Cambridge, Cambridge, United Kingdom; Smithsonian Tropical Research Institute, Panama City, Panama; Centre for Ecology and Conservation, College of Life and Environmental Sciences, University of Exeter, Penryn, United Kingdom

**Keywords:** aposematism, light environment, avian vision, butterfly vision, colour signal

## Abstract

Forests are a mosaic of light spectra, and colour signal efficiency might change in different light environments. Local adaptation in *Heliconius* butterflies is linked to microhabitat use and the colourful wing colour patterns may be adapted for signalling in different light environments. These toxic butterflies exhibit conspicuous colours as a warning to predators that they should be avoided, but also find and choose potential mates based on colour signals. The two selection pressures of predation and mate preference are therefore acting together. Colour conspicuousness should show habitat-specific contrast for the butterflies, which would facilitate detection and species identification. On the other hand, selection for signal stability would be stronger in the avian visual system. In this study we analysed the contrast of two *Heliconius* mimicry rings in their natural habitats under varying degrees of forest fragmentation and light conditions. We used digital image analyses and mapped the bird and butterfly vision colour space in order to examine whether warning colours have greater contrast and if they transmit a consistent signal across time of the day and habitat in a tropical forest. We tested conspicuousness using opponent colour channels against a natural green background. For avian vision, colours are generally very stable through time and habitat. For butterfly vision, there is some evidence that species are more contrasting in their own habitats, where conspicuousness is higher for red and yellow bands in the border and for white in the forest. Light environment affects *Heliconius* butterflies’ warning signal transmission to a higher degree through their own vision, but to a lesser degree through avian predator vision. This work provides insight into the use of colour signals in sexual and natural selection in the light of ecological adaptation.

## Introduction

The success of a signal is related to its effectiveness in a specific environment and how strongly it influences the behaviour of the receiver (Endler 1978). Forests are a mosaic of light colours, and the same colour pattern can have an altered appearance in different light environments (Endler 1993). If an individual shows high reflectance of a specific wavelength, but the environment lacks light in that part of the spectrum, the region of high reflection will be unimportant as a signal (Stevens et al. 2007). Ambient light spectra also vary from dawn to dusk, hence species that signal only at certain times and places are expected to evolve characteristics and predictable combinations of colours for particular environments (Endler 1993). Therefore, ambient light characteristics should be included together with the receiver visual system to understand the microhabitat choice and behaviour of animals.

In the case of colour signals, perception can be altered by the habitat they live in by filtering wavelengths and altering visual backgrounds (Endler 1993; Lovell et al. 2005). Sensory drive explains the process of adaptation of signalling and sensory systems to the local environment (Endler 1992; Endler and Basolo 1998). Environment tuned spectral sensitivity is better known in aquatic habitats, such as in guppies (Endler 1980) and cichlid fish (Seehausen et al. 2008), as compared to terrestrial light environments. On land, colour depends on the reflection of the surroundings and has greater variability over time (Boughman 2002). Habitat signal transmission can favour diversification of mating signals through local adaptation, leading to reproductive isolation. Distinct *Anolis* lizards male dewlaps are found in different microhabitats (Fleishman et al. 1997). Male dewlap colours are more conspicuous in their own habitat than in other habitats, mainly because of the contrast against the background in the ultraviolet (UV) range (Leal and Fleishman 2002). Perception of colours in different light conditions can also influence attacks by predators, for example among butterflies in an environment with high UV light, birds aimed at the butterfly wings, more specifically the marginal white eyespots that have UV reflectance, instead of the head (Olofsson et al. 2010).

Local adaptation in *Heliconius* butterflies commonly involves adaptation to specific microhabitat use (Estrada and Jiggins 2002; Elias et al. 2008; Jiggins 2008). Mimicry rings are groups of unpalatable species that share the same warning colour, and these tend to be found in specific microhabitats such as forest or open areas. The *Heliconius* habitats are associated with the use of larval host-plants, adult food plants, sexual behaviour and gregarious roosting (Mallet and Gilbert 1995). Species that lay eggs on *Passiflora* species that occur in second growth tend to be seen in open areas, while species that lay eggs on canopy *Passiflora* vines are seen flying high in the forest. The choice of microhabitat also might be connected with light differences between those environments, such as the choice of using shady areas in communal roosting (Mallet and Gilbert 1995; Finkbeiner 2014).

Therefore, different light environments should create microhabitats where butterfly signals would be more efficient. Although mimicry rings differ in their microhabitat, the light environment has not been measured to verify whether colour patterns could be specifically adapted to particular light environments. The colourful wing colours of *Heliconius* butterflies may also be subject to evolution caused by sensory drive due to their potentially conflicting roles in predation and mate preference. Many species exhibit Müllerian mimicry (Müller 1879), in which two or more species share the same conspicuous colour as a warning to predators that they are toxic and should be avoided (Benson 1972). Also, these butterflies find and choose potential mates based on colour signals, which can lead to reproductive isolation (Jiggins et al. 2001; Sweeney et al. 2003; Kronforst et al. 2006). Furthermore, communication between conspecifics might be based on UV signals, since *H. erato* females express the duplicate UV opsin gene, which allows a greater degree of discrimination of the UV-yellow wing patches (Briscoe et al. 2010; Bybee et al. 2012; McCulloch et al. 2016; Dell’Aglio et al. 2018).

This microhabitat structuring allows mimicry rings to remain distinct. This may be because there are sets of predators in different habitats, each of which perceive a different mimicry ring as the most abundant pattern (Joron and Mallet 1998). Although little is known of the specific predators that attack *Heliconius*, it seems likely that their aposematic signals are directed at several predators with different visual abilities and spectral sensitivities (Dell’Aglio et al. 2018). Ambient light together with predator sensitivity can interfere with the interpretation of the information perceived from colour signals.

Warning coloration should, therefore, be easy to detect and memorize even in heterogeneous environments and light conditions (Guilford and Dawkins 1991; Endler 1992). Warning signals are often dominated by red, yellow and orange, frequently contrasting with black, which are the main colours in *Heliconius*. The reason why these long-wavelength colours are widely represented in aposematic coloration is that they are highly conspicuous against natural backgrounds, are more stable across light conditions, allowing long distance discrimination and detectability, and influence memorability (Guilford and Dawkins 1991; Stevens and Ruxton 2012; Arenas et al. 2014; Dell’Aglio et al. 2016).

Perception of colour depends on several neurophysiological mechanisms, such as the presence of opponent colour channels. This chromatic mechanism involves comparisons of receptors outputs, in which opposite neural pathways are either activated or inhibited depending on the stimuli reaching the eye (Kelber et al. 2003; Renoult et al. 2015). This mechanism is useful especially regarding colour stability against spatial and temporal variation in illumination (Lovell et al. 2005; Renoult et al. 2015). For example birds, the major predator of aposematic butterflies, have tetrachromatic vision and seemingly have at least three opponent channels, as found in domestic chicks (Vorobyev et al. 1998; Osorio et al. 1999). Opponent channels have also been described for insects (Chittka et al. 1992; Chittka 1996) and butterflies (Kelber 1999), and have been hypothesized for *Heliconius* butterflies although more behavioural analyses are needed to confirm which opponent channels are actually used (Swihart 1971, 1972; Bybee et al. 2012). Butterflies in the genus *Papilio* have duplicate LW opsin genes to see in the red and green range (Kelber 1999; Briscoe 2008), while *Heliconius* has differences in the red and green photoreceptor sensitivity associated with the presence of red filtering pigments in the ommatidia (Zaccardi et al. 2006; McCulloch et al. 2016). Thus, we expect avian predators and butterflies to rely on these high-contrast systems to process information under a changing light environment.

The aim of this study was to analyse *Heliconius* warning colouration under different light conditions in their natural habitats. In particular, to test conspicuousness of the wing colouration against a natural green background, encoded by opponent colour channels. Using digital image analyses, butterfly wings were photographed and mapped to UVS and VS avian predator vision and to *Heliconius erato* vision (Figure 1). Our predictions are that (1) signal contrast and conspicuousness for avian predators should have stability, that should be reliable throughout the day and in different light environments (Stevens and Ruxton 2012; Arenas et al. 2014). Warning signals might be honest indicators of prey unprofitability to predators, and if signals fluctuate through the day and between light environments, we would predict that this could delay learning by predators and be costly to the prey. Similarly for internal contrasts (i.e. contrast between black and the coloured bands), therefore conspicuousness would not rely totally on background contrast but also on internal patterns which account for close-distance conspicuousness (Endler 1978; Aronsson and Gamberale-Stille 2009). (2) From a *Heliconius* butterfly perspective; we predict that signal contrast and conspicuousness should show habitat-specific maximum background contrast and higher colour differences in their own habitats (Table 1), which would facilitate detection and species identification. We therefore predict that selection for signal stability will be much stronger in the avian visual system as compared to the butterfly visual system.

**Figure 1.**
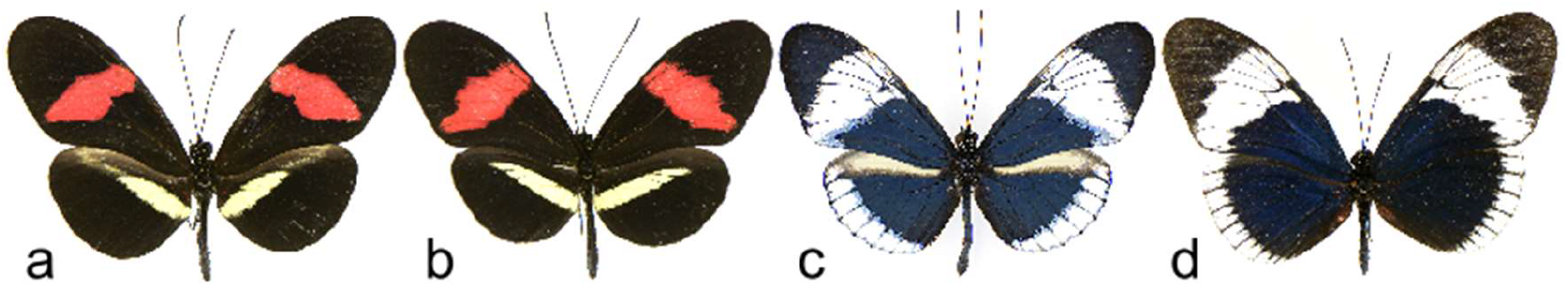
Co-mimics used in this study. (a) *Heliconius erato demophoon*. (b) *H. melpomene rosina*. (c) *H. cydno*. (d) *H. sapho*.

**Table 1.** Colour patches for each co-mimic pair studied and typical microhabitats and light conditions where these co-mimics occur. Microhabitats descriptions are based on Estrada and Jiggins (2002) and light conditions based on Endler (1993) and personal observations.

| Co-mimics | Colour | Microhabitat | Light conditions |
| --- | --- | --- | --- |
| *H. erato demophoon* & *H. melpomene rosina* | red & yellow | forest border | partial shade |
|  |  | open area | no shade |
| *H. sapho* & *H. cydno* | white | closed forest | full shade |
|  |  | canopy | partial shade |

## Material and Methods

### Study site and species

Fieldwork was performed during the dry season, along Pipeline Road, a tropical lowland rainforest in the Panama Canal Zone (Parque Nacional Soberanía, 9°7’33”N, 79°42’90”W). Pipeline Road makes a transect through the forest, creating a heterogeneous habitat with open sunny areas and close canopy exceeding 30 m in height. All specimens were collected in the area. Two pairs of co-mimics that live in sympatry were selected, *H. erato demophoon* (n = 8) and *H. melpomene rosina* (n = 8), *H. sapho* (n = 5) and *H. cydno* (n = 5), each pair belonging to a different mimicry ring, red and yellow, and white, respectively (Figure 1 and Table 1).

### Digital photography

The general approach and methodology for this work was based on previous work with colour stability using opponent signals (Lovell et al. 2005; Arenas et al. 2014). The spectral reflectance of mimetic pairs was investigated using digital photography. This provides a way to control for natural variation in luminance intensity (shadowing) that is not captured by spectrometry, and also allows non-invasive colour measurements easily applied in the field (Stevens et al. 2007; Troscianko and Stevens 2015). Therefore, through this method we could obtain colour measures under the sensitivity of all receiver photoreceptors (300-700 nm) in the actual viewing conditions of conspecifics and avian predators.

Fresh wings of each specimen were photographed following the same methods of image collection in Dell’Aglio et al. (2018). The camera was fitted to a tripod and pointed towards the ground (90°) at a height of approximately 80 cm. Each photo setup included two individuals, one of each species of the co-mimic pair, a 40% grey standard (Spectralon® Labsphere) used for calibration and a leaf freshly collected to make background measures. The species used was *Guazuma ulmifolia* (Sterculiaceae), a small abundant shrub across all Pipeline Road, which facilitated the collection of fresh leaves.

Photos were taken under three different arboreal canopy conditions, forest border, closed forest, and open area, where those butterflies are usually seen (Table 1). All photos were taken under sunny to part-cloudy days, with three replicates in each habitat making sure that the amount of light was similar. In order to standardize the replicates, light measures were taken with a digital light meter (Digital Lux meter, Tondaj LX-1010B), which measures the total amount of LW (555 nm) per square meter (Lux) (Figure 2). Also photographs of the canopy were taken in order to measure vegetative cover, which was 81.5% (SE ± 0.2) for closed forest, 59.8 % (SE ± 3.4) for forest border, and 0% for open area. The aim was to analyze how colour signals are perceived throughout the morning when butterflies are most active. Therefore, photos were taken at dawn (7 am), morning (9 am) and noon (12 pm) during a short period of 15min as light conditions change rapidly. Open area photographs were taken only at 12 pm to represent a highly used environment by butterflies at this period of day.

**Figure 2.**
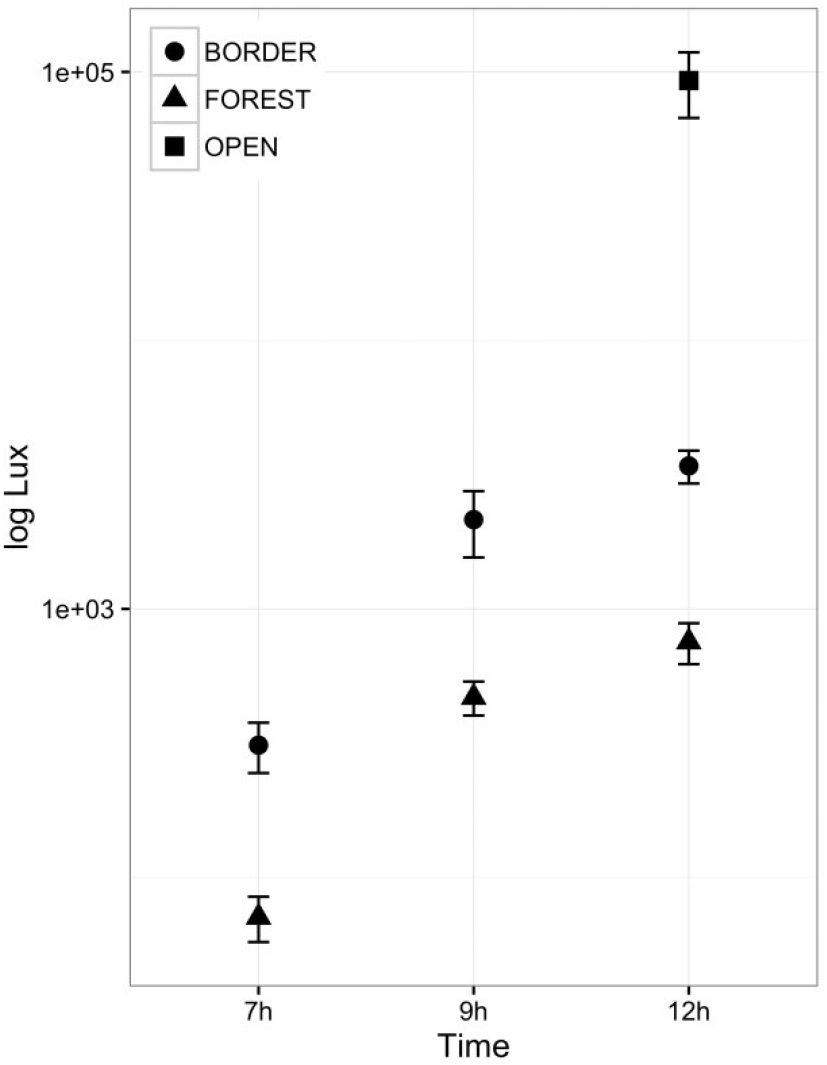
Amount of light differs across light environment and time of day. Showing the average log Lux between the three replicates of each habitat (± SE). Lux represents the amount of long-wave (555 nm) per square meter.

### Image analysis and visual modelling

All images were processed and analysed into the imaging software ImageJ (Rasband, 1997-2012). RAW human-visible and UV images were linearized and aligned following the methodology of Troscianko and Stevens (2015) and Arenas et al. (2014). Normally, photos would be normalized to the grey standard, which removes effects of light conditions (Arenas et al. 2014). Since our main interest was to measure how coloration changes in different light environments, the images were not normalized. Instead, an average grey standard value was obtained from all photographs. Photon catch values were obtained for each colour using the entire patch from linearized photos, and subsequently these values were multiplied by each photo exposure time and normalized with the average grey standard. With this methodology we were able to calculate how particular environment and time varies from the natural average light (Stevens et al. 2007; Arenas et al. 2014). We used the average photon catch results from the three habitats replicates. Predicted photon catch values were obtained using spectral sensitivity for each cone type of the blue tit (*Cyanistes caeruleus*) for the UV-sensitive vision (UVS) (Hart et al. 2000), peafowl (*Pavo cristatus*) for the violet-sensitive vision (VS) (Hart 2002) and the female *Heliconius erato* (Briscoe et al. 2010; McCulloch et al. 2016).

Background of many terrestrial habitats is dominated by greenish vegetation; therefore, a green leaf was chosen to make contrast calculations. Differences between light environment and time of the day were calculated using the contrast of warning colours against an average green leaf. Channel activation in avian vision was calculated using the Red-Green (RG), Blue-Yellow (BY) and Blue-UV opponent channel (Osorio et al. 1999; Lovell et al. 2005; Stevens et al. 2009). For the achromatic signal, we used avian double cones (DBL). Using a ratio-based approach suggested by Lovell et al. (2005), we calculated the opponent channel responses as follows:

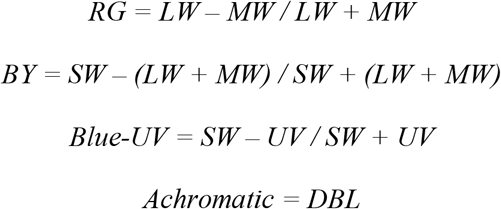

The *Heliconius* opponent channels were calculated based on what has been proposed in earlier studies regarding *Heliconius* vision (Swihart 1971, 1972; Bybee et al. 2012). The *H. erato* compound eye has red filtering pigments that shift LW photoreceptor sensitivity from green to red and as the physiological mechanisms underlying these two LW photoreceptors are not known, sensitivities in Green (560 nm) and Red (600 nm) were used (McCulloch et al. 2016, 2017). To investigate differences between co-mimic species, we calculated opponent channel activation based on the prediction that *Heliconius* mating system might use UV and Red/Green light for mate choice (Bybee et al. 2012; McCulloch et al. 2016, 2017). Opponent channels in *Heliconius* vision was calculated using as follows:

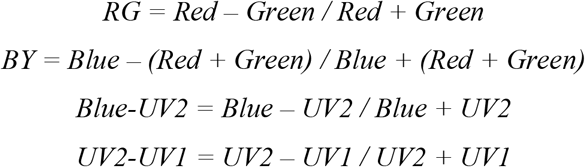

To examine whether warning colours have greater contrast against green background we calculated the Weber Contrast (Whittle 1994), which takes into account the image value of the objects of interest as a fraction of background appearance using the formula:

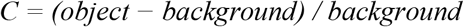

Where *background* corresponds to the green leaf opponent channel values, and *object* corresponds to warning colour opponent channel values. This measure is suited to comparisons between small objects against larger backgrounds, such as butterflies against the green forest. For internal contrast, achromatic values of the warning colours were used against the black of each individual wing as background (Arenas et al. 2014). We plotted the mean absolute contrast of each colour signal as a function of time and light environment for the three vision models.

To test contrast stability through time and habitat, we used the coefficient of variation (CV) of each colour in each opponent channel for each visual system. The CV is an effective measurement to determine how relatively stable a measurement is around a mean value. The CV can be interpreted as greater its value, more variance and instability the signal would be, following methodology from Arenas et al. (2014).

### Statistical analyses

All statistical calculations were processed in the software R 3.5.3 (R Core Team 2020). Our approach was to model colour contrasts over the course of a day and under different habitats in term of both predator and butterfly vision. General linear mixed-effects models were performed using random effects (package *lme4*) followed by Tukey’s post-hoc (package *multicomp*). Normality tests showed that contrast data were not normally distributed, therefore *glmer* tests were performed using *Gama* distribution. Raw data was plotted to illustrate the results (package *ggplot2*). The models were fitted accordingly to the predictions outlined above. Analyses were carried out using contrast values as the dependent variable, and fixed and random factors varied depending on the question. Factors were individuals, colour (red, yellow, white), habitat (border, forest, area), time (7am, 9am, 12pm), and bird vision (UVS, VS). For *Heliconius* vision, we also added side of the wing (dorsal, ventral).

## Results

### Signal contrast and conspicuousness for avian predators

Red was generally the most contrasting colour against a green background in the RG opponent channel as compared to yellow (z = 14.05, *P* < 0.001, Table S1) and white (z = 17.68, *P* < 0.001, Table S1). In contrast, white had higher contrasts against a green background in the BY channel, as compared to red (z = −15.3, *P* < 0.001, Table S1) and yellow (z = 22.04, *P* < 0.001, Table S1) (Figure 3). Colours in open areas showed a higher contrast, such in the RG channel for red band (t = −6.16, *P* < 0.001, Table S2) with no difference between border and forest (z = −0.31, *P* = 0.94, Table S2) (Figure 3). In the Blue-UV opponent channel, UVS and VS birds could perceive red and yellow with less stability (Table 2), as red and yellow showed higher contrast in the morning than at 12 pm (Table S3). On the other hand, white colour in the Blue-UV channel did not show any changes through the day (Table S3).

**Figure 3.**
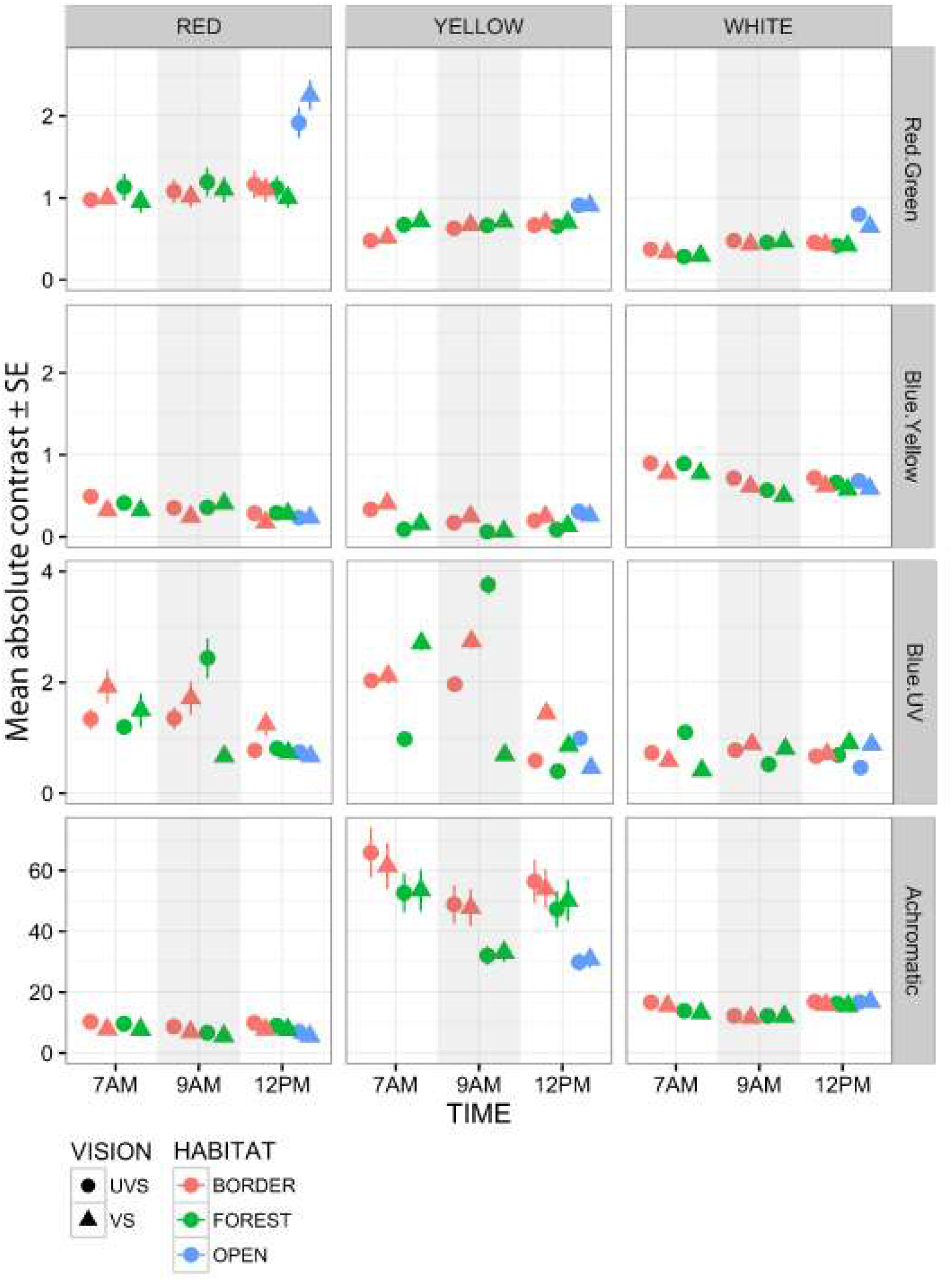
Colour conspicuousness for avian predators. Mean absolute contrast of colour signals (±SE, standard error) in the bird vision systems analyzed (circles, UVS; triangles, VS) through habitats (red, border; green, forest; blue, open) and time (7am, 9am, 12pm). Vertical panels show the three colour signals (red, yellow and white), horizontal panels show opponent channels against green leaf (top, Red-Green; middle, Blue-Yellow and Blue-UV) and against the black of the wing (bottom, Achromatic). Note: Channels have different y-axis values. Error bars smaller than data points are not shown.

**Table 2.** *Heliconius* vision has higher coefficient of variation in most of the colours compared with bird vision across time and habitats. Coefficient of variation for each visual system, colour and opponent channel, higher values are in bold.

| Opponent channel | Colour | Visual system |  |  |
| --- | --- | --- | --- | --- |
|  |  | *Heliconius* | UVS | VS |
| Red-Green |  |  |  |  |
|  | Red | **1.053** | 0.704 | 0.695 |
|  | Yellow | **0.384** | 0.211 | 0.178 |
|  | White | **0.722** | 0.375 | 0.314 |
| Blue-Yellow |  |  |  |  |
|  | Red | **0.699** | 0.576 | 0.606 |
|  | Yellow | **0.735** | 0.667 | 0.528 |
|  | White | **0.177** | 0.134 | 0.169 |
| Blue-UV |  |  |  |  |
|  | Red | 0.783 | 0.897 | **1.024** |
|  | Yellow | 0.265 | **0.788** | 0.637 |
|  | White | **1.035** | 0.549 | 0.341 |
| UV2-UV1 |  |  |  |  |
|  | Red | 0.525 | - | - |
|  | Yellow | 1.047 | - | - |
|  | White | 1.061 | - | - |
| Achromatic |  |  |  |  |
|  | Red | - | 0.367 | 0.436 |
|  | Yellow | - | 0.717 | 0.699 |
|  | White | - | 0.348 | 0.349 |

Internal achromatic contrast was higher for yellow, compared to red (z = −36.7, *P* < 0.001, Table S1) and white (z = −22.9, *P* < 0.001, Table S1). Moreover, yellow had more contrast in the border (Figure 3), which is the preferred habitat of yellow band butterflies, than in the forest (z = 3.5, *P* = 0.001, Table S4).

### Signal contrast and conspicuousness for Heliconius conspecifics

In some cases, contrasts followed our prediction that species would be more contrasting in their own habitats (Figure 4). The yellow colour was more contrasting in the border in the UV2-UV1 channel, especially early hours such as 7 am (t = 6.9, *P* < 0.001, Table S5) and 9 am (t = 354.3, *P* < 0.001, Table S5), but not at 12 pm (t = 2.2, *P* = 0.218, Table S5). In the Blue-UV2 channel, white was more contrasting in the forest than in the border at 7 am (t = −3.9, *P* < 0.001, Table S5). Although, while white colour contrast decreased during the day in the forest, it increased at 12 pm in the border (t = 4.77, *P* < 0.001, Table S5) (Figure 4).

**Figure 4.**
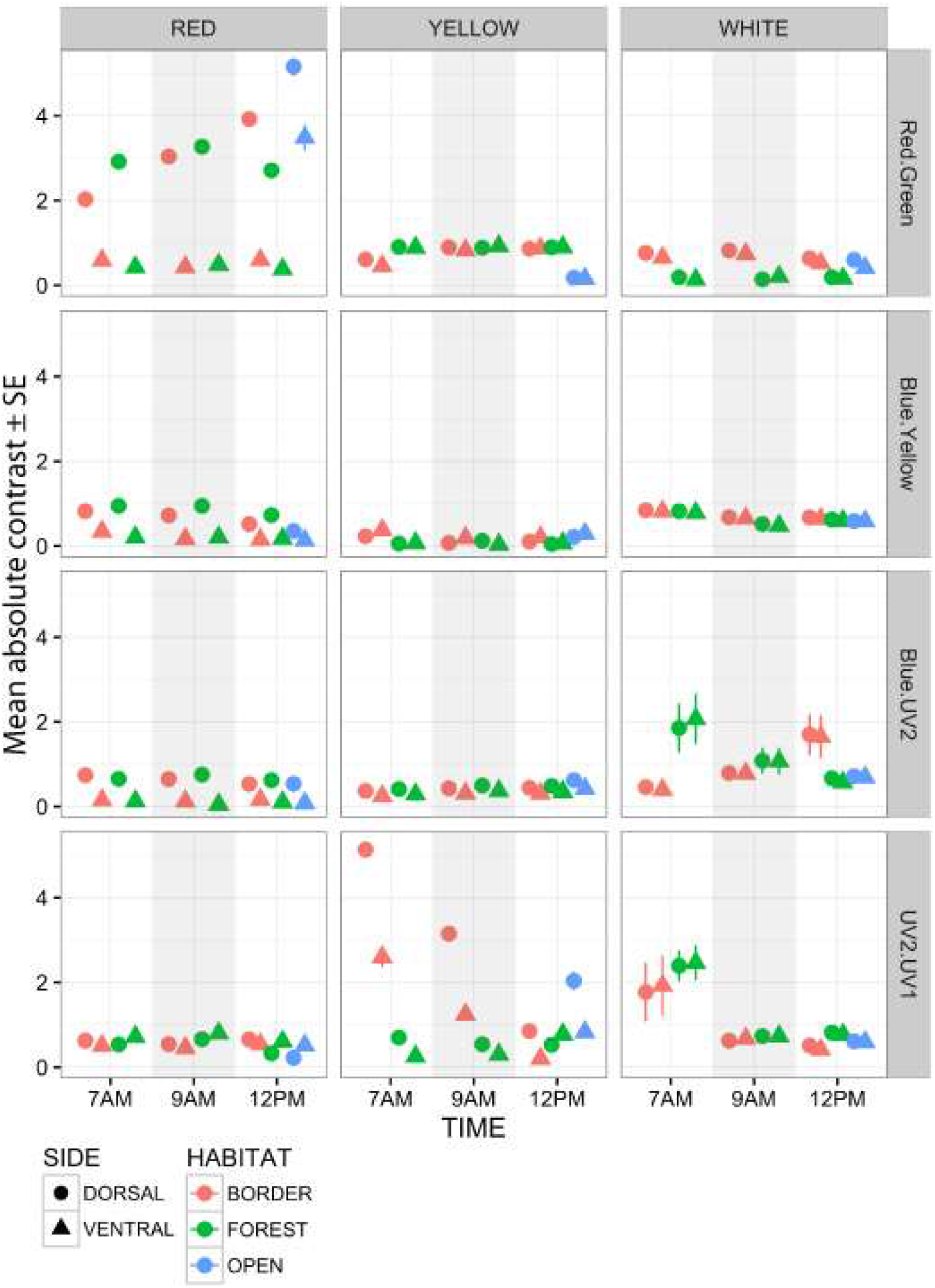
Colour conspicuousness for *Heliconius* conspecifics. Mean absolute contrast of colour signals against green leaf in *Heliconius* vision through habitats (red, border; green, forest; blue, open), time (7 am, 9 am, 12 pm) and side of the wing (circles, dorsal; triangles, ventral). Vertical panels show colour signals (red, yellow and white), horizontal panels show opponent channels against green leaf (top, Red-Green; middle, Blue-Yellow and Blue-UV2; bottom, UV2-UV1). Error bars: ± 1 standard error (SE), error bars smaller than data points are not shown.

The red colour showed large differences in the RG channel between dorsal and ventral side, with dorsal side with the higher contrast (t = 10.1, *P* < 0.001, Table S6). Same results were found for red in the Blue-Yellow and Blue-UV2 opponent channel (Table S6) (Figure 4).

The contrast of signals against the background revealed greater differences across habitat and time when seen through the *Heliconius* vision model (Table 2). *Heliconius* vision had higher values of coefficient of variation, which revealed greater fluctuations across habitat and time in most of the opponent channels. The only exception was in the Blue-UV opponent channel for red and yellow colour, which showed more instability for avian vision than for *Heliconius* vision (Table 2).

## Discussion

The bright and contrasting *Heliconius* wing patterns appear well adapted for signalling distastefulness to predators. However, their colour constancy and appearance in different light environments remains poorly studied. Here we have shown that colours are indeed very stable for avian predator vision, but somewhat less so for *Heliconius* vision. This is consistent with the idea that wing patterns are primarily selected for their role in signalling distastefulness to predators.

### Signal stability and conspicuousness to avian predators

Generally, warning signals involve combinations of long-wavelength colours such as red, orange, and yellow which are highly conspicuous against natural backgrounds and stable under different natural conditions (Lovell et al. 2005; Stevens and Ruxton 2012). In our results, *Heliconius* red colouration has higher detectability against average green background in the RG output and these results are consistent regardless of habitat and time of the day. Previous work has investigated colour stability through opponent colour channels and also showed that red coloration is more contrasting and stable against green backgrounds over the course of a day and across light conditions to the bird visual system (Lovell et al. 2005; Arenas et al. 2014). Moreover, the *Heliconius* yellow colouration also is highly conspicuous against its internal black pattern in the achromatic output. Achromatic information is one of the main cues used for motion detection (Hämäläinen et al. 2015). Therefore, our results suggest that red and yellow signals work together and are likely effective in stimulating avian opponent channels in order to be conspicuous in all light environments.

There is evidence that red and yellow colouration serve as reliable warning signal to avian predators (Ham et al. 2006; Svádová et al. 2009; Arenas et al. 2015), but that this is less true of white colouration. One explanation is that white is more variable across time and habitat, so provides a less reliable signal under varied light conditions (Stevens and Ruxton 2012; Arenas et al. 2014). As an example, field and aviary experiments with polymorphic yellow and white wood tiger moths, *Parasemia plantaginis*, showed that yellow males are avoided more than white males by predators, but white males have higher mating success (Nokelainen et al. 2012). Our results showed that white contrasts against green background were lower and rather variable for avian vision. The co-mimics *H. sapho* and *H. cydno* also contain iridescence blue that was not measured with this methodology. However, the lack of high contrast in white colouration might be balanced with the fact that polarized light might act as a signal, especially in forest habitats (Sweeney et al. 2003; Douglas et al. 2007; Pegram et al. 2015).

Highly conspicuous warning signals are expected to evolve to be stable in their appearance throughout the day and between light environments, in order to remain honest indicators of prey unpalatability (Blount et al. 2009; Cortesi and Cheney 2010; Stevens and Ruxton 2012; Arenas et al. 2015). If warning signals fluctuate through time and space this could alter bird foraging experiences and reduce the effectiveness of the aposematic signal. The final decision on whether or not to attack a prey results from a combination of information reaching the predator brain, and for greater efficiency, aposematic coloration needs to be easy to remember (Endler 1988). Our results support this prediction, as colours were generally stable through time and light environments in all opponent systems with only a few exceptions. Notably these occurred where contrasts were higher in open areas and in the early morning. This might also be favourable as the prey would be more conspicuous when they are most vulnerable to predation, since birds are more active and forage early in the morning (Buskirk et al. 1972; Poulin et al. 2001; Steiger et al. 2009). In agreement with this, *Heliconius* predation and roost disturbance has been observed in the early morning (Mallet 1986; Finkbeiner 2014).

### Habitat and time influence conspicuousness in Heliconius conspecifics

Butterflies belonging to the two mimetic rings studied here tend to be segregated between habitats, corresponding to areas where the photographs were taken, although there is considerable overlap (Estrada and Jiggins 2002). We showed that the colours were more unstable when seen through *Heliconius* vision as compared to avian vision and some colours tend to be more contrasting in their respective habitats.

Our results provide some evidence that co-mimic rings are more conspicuous in their own habitat as seen through *Heliconius* vision, reinforcing the idea that ecological adaptation leads to spatial segregation to where detection would be facilitated. Some colours had higher contrast against green backgrounds in their respective habitat, such as yellow in the border and white in the forest. Nonetheless, red showed the opposite trend and was generally more contrasting in the forest. Differences across light environments could affect mating preferences by altering search costs for a specific colour pattern, and perhaps changing the fitness of different colour patterns. Adaptation in different microhabitats within the forest might have an influence on how closely related species commonly differ in pattern, while convergence in pattern occurs between more distantly related species (Joron and Mallet 1998). Ecological adaptation is attributed to habitat preference and leads to assortative mating (Jiggins 2008). The two sister species studied here, *H. melpomene rosina* and *H. cydno*, are known to rarely hybridise in the wild, hence microhabitat segregation reduces potential mating encounters between these two species and reduces gene flow (Mallet et al. 1998; Merrill et al. 2013). Subtle environmental conditions could affect recognition in mating behaviour as seen in the jumping spider, *Habronattus pyrrithrix*, which red males were more successful in approaching females in the sunlight (Taylor and McGraw 2013).

The activation of opponent channels was often higher in the early hours of the morning, at the time when the butterflies are more active and leave their roost or perches to forage (Mallet 1986; Finkbeiner et al. 2012). This was especially the case for *Heliconius* white and yellow wing colours in the UV channel, which might act in intraspecific communication (Briscoe et al. 2010; Bybee et al. 2012). There is evidence that distinct UV colour signals are being transmitted between co-mimics, which may reduce costs of mating confusion (Dell’Aglio et al. 2018). Similarly, in two species of newt, belly colour is distinct in the UV range and females often made mistakes choosing the wrong males in the absence of UV light (Secondi and Théry 2014).

In this context, the duplicate genes encoding two distinct visual pigments with sensitivity peaks in the UV range in *H. erato* females offer the potential for enhanced spectral discrimination in light environments and time of the day where UV is more prominent. A UV2-UV1 opponent channel was proposed by Bybee et al. (2012), who showed that this receptor combination would have lower error rates for discrimination between *Heliconius* and *Dryas* yellows. There is no direct evidence for such a mechanism yet, but the fact that males and females show differences in the expression of the two UV proteins suggests that UV2-UV1 contrasts could be an important opponent channel for a female specific behaviour, perhaps mate recognition or host plant finding (McCulloch et al. 2016, 2017). On the other hand, there is some evidence for a Red-Green opponent channel seen in behavioural trials, which showed that *Heliconius erato* use both green and red photoreceptor to detect colours in the 590-640 nm range (Zaccardi et al. 2006) and the presence of red filters in their eyes means that this is a possibility (McCulloch et al. 2016). Differences between species in Red-Green channel activity for red colouration might have a role in mate recognition since *Heliconius* tend to be attracted to red (Merrill et al. 2011).

The sensory drive hypothesis describes evolutionary relationships among visual systems, conditions of the light environment and mating preferences (Endler and Basolo 1998). *Heliconius* mating preference is highly linked to colour and in *H. melpomene*, the gene responsible for red colour pattern is genetically linked to the preference for the same pattern (Jiggins et al. 2001; Naisbit et al. 2001; Merrill et al. 2011). Visual sensitivity data used here is only from *H. erato* females whereas it might differ for other species in their visual systems and perhaps match with colour preference or habitat (Frentiu et al. 2007; Briscoe et al. 2010; McCulloch et al. 2017). In addition, mating behaviour might benefit from some habitats in maximizing conspicuousness, such as in tropical dwelling birds and wire-tailed manakins which visual contrast is increased during display by habitat choice (Endler and Théry 1996; Heindl and Winkler 2003). Nevertheless, our results suggest that selection for conspicuousness in the preferred habitat could explain in part the divergence in colour pattern in these species.

### Conclusion

In conclusion, the transmission of *Heliconius* warning signals varies due to light environment to a much greater degree through their own visual system, but to a smaller degree through avian predator vision. Selection for signal detectability under different habitat conditions is a mechanism that is proposed to lead to evolution of signal diversity, as seen in species of *Anolis* lizards that occupy habitats that match their visual system and signal design (Leal and Fleishman 2002), in species of warblers which different cone opsin gene expression correlate with sexual selection and habitat use (Bloch 2015) and also colour patterns of guppies are more conspicuous to guppies at the times and places of courtship and relatively less conspicuous at times and places of predator risk (Endler 1991). *Heliconius* butterfly warning colours are highly contrasting against the forest background and stable through time and habitat in terms of predator avoidance but also conspicuous to attract the attention of conspecifics. However, more extensive studies considering spectral sensitivities of different *Heliconius* species and their responses to environmental changes in their signal visibility are needed to confirm the conspicuousness to mates. Opponent channel colour contrasts can predict behaviour of perceivers, however, additional behavioural experiments on how light environment influences prey detectability, such with poison frogs (Rojas et al. 2014), are necessary to verify our results.

## Author contributions

D.D.D. collected all the data, analysed, and wrote the manuscript; J.T. developed digital image methodology and data analysis; W.O.M, M.S, and C.D.J conceived the study, edited, and wrote the manuscript. We have no conflict of interest to declare.

## Supplementary Information

**Table S1.**
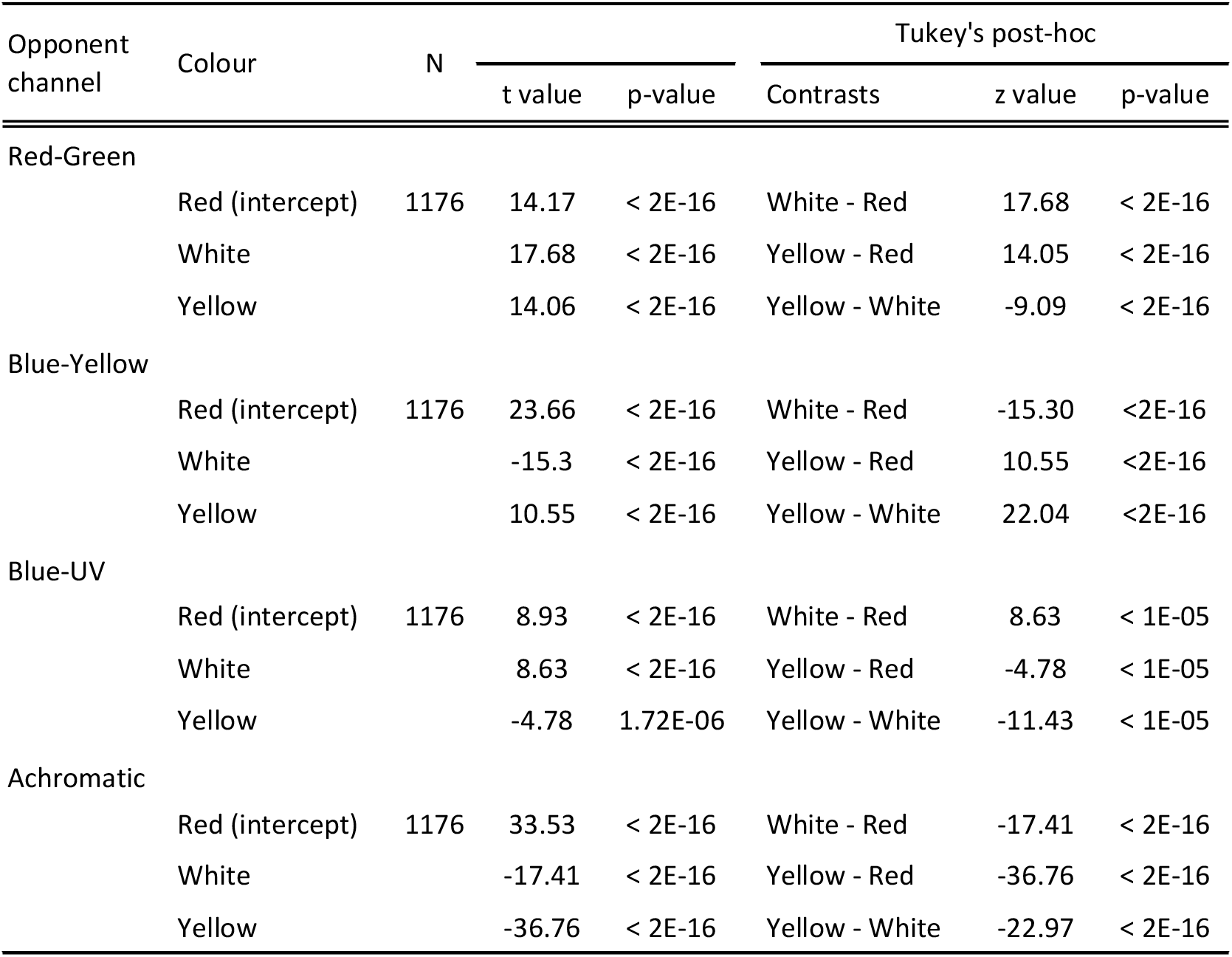
Colour contrast differences for avian vision per opponent channel. Generalized linear mixed-effects models with random effects and Tukey’s post-hoc (y ~ Colour + (1 | Individuals) + (1 | Vision: Habitat: Time)).

**Table S2.** Habitat contrast differences for avian vision per colour for the Red-Green opponent channel. Generalized linear mixed-effects models with random effects and Tukey’s post-hoc (y ~ Habitat + (1 | Individuals) + (1 | Vision:Time)).

| Colour | Habitat | N | Linear mixed model |  | Tukey's post-hoc |  |  |
| --- | --- | --- | --- | --- | --- | --- | --- |
|  |  |  | t value | p-value | Contrasts | z value | p-value |
| Red |  |  |  |  |  |  |  |
|  | Border (intercept) | 448 | 16.59 | <2E-16 | Forest - Border | -0.31 | 0.947 |
|  | Forest |  | -0.31 | 0.753 | Open - Border | -6.16 | <1E-05 |
|  | Open |  | -6.16 | 7.28E-10 | Open - Forest | -5.91 | <1E-05 |
| Yellow |  |  |  |  |  |  |  |
|  | Border (intercept) | 448 | 23.56 | <2E-16 | Forest - Border | -8.46 | <2E-16 |
|  | Forest |  | -8.46 | <2E-16 | Open - Border | -16.69 | <2E-16 |
|  | Open |  | -16.69 | <2E-16 | Open - Forest | -10.65 | <2E-16 |
| White |  |  |  |  |  |  |  |
|  | Border (intercept) | 280 | 9.07 | <2E-16 | Forest - Border | 2.76 | 0.015 |
|  | Forest |  | 2.76 | 0.006 | Open - Border | -12.39 | 0.001 |
|  | Open |  | -12.39 | <2E-16 | Open - Forest | -14.39 | 0.001 |

**Table S3.** Time contrast differences for avian vision per colour for the Blue-Yellow opponent channel. Generalized linear mixed-effects models with random effects and Tukey’s post-hoc (y ~ Time + (1 | Individuals) + (1 | Vision:Habitat)).

| Colour | Time | N | Linear mixed model |  | Tukey's post-hoc |  |  |
| --- | --- | --- | --- | --- | --- | --- | --- |
|  |  |  | t value | p-value | Contrasts | z value | p-value |
| Red |  |  |  |  |  |  |  |
|  | 12PM (intercept) | 448 | 12.73 | <2E-16 | 7AM - 12PM | -5.11 | <1E-06 |
|  | 7AM |  | -5.11 | 3.2E-07 | 9AM - 12PM | -5.39 | <1E-06 |
|  | 9AM |  | -5.39 | 6.7E-08 | 9AM - 7AM | -0.33 | 0.939 |
| Yellow |  |  |  |  |  |  |  |
|  | 12PM (intercept) | 448 | 8.94 | <2E-16 | 7AM - 12PM | -10.38 | <1E-04 |
|  | 7AM |  | -10.38 | <2E-16 | 9AM - 12PM | -11.64 | <1E-04 |
|  | 9AM |  | -11.64 | <2E-16 | 9AM - 7AM | -2.05 | 0.092 |
| White |  |  |  |  |  |  |  |
|  | 12PM (intercept) | 280 | 10.6 | <2E-16 | 7AM - 12PM | 0.4 | 0.909 |
|  | 7AM |  | 0.41 | 0.677 | 9AM - 12PM | -0.23 | 0.970 |
|  | 9AM |  | -0.23 | 0.813 | 9AM - 7AM | -0.64 | 0.797 |

**Table S4.** Habitat contrast differences for avian vision for the achromatic opponent channel per colour. Generalized linear mixed-effects models with random effects and Tukey’s post-hoc (y ~ Habitat + (1 | Individuals) + (1 | Vision:Time)).

| Colour | Habitat | N | Linear mixed model |  | Tukey's post-hoc |  |  |
| --- | --- | --- | --- | --- | --- | --- | --- |
|  |  |  | t value | p-value | Contrasts | z value | p-value |
| Red |  |  |  |  |  |  |  |
|  | Border (intercept) | 448 | 8.19 | 2.49E-16 | Forest - Border | 2.94 | 0.008 |
|  | Forest |  | 2.94 | 0.003 | Open - Border | 6.30 | 0.001 |
|  | Open |  | 6.30 | 2.93E-10 | Open - Forest | 4.64 | 0.001 |
| Yellow |  |  |  |  |  |  |  |
|  | Border (intercept) | 448 | 8.85 | <2E-16 | Forest - Border | 3.52 | 0.001 |
|  | Forest |  | 3.52 | 4.3E-04 | Open - Border | 5.75 | 0.001 |
|  | Open |  | 5.75 | 8.5E-09 | Open - Forest | 4.07 | 0.001 |
| White |  |  |  |  |  |  |  |
|  | Border (intercept) | 280 | 20.06 | <2E-16 | Forest - Border | 2.12 | 0.083 |
|  | Forest |  | 2.12 | 0.034 | Open - Border | -0.33 | 0.940 |
|  | Open |  | -0.33 | 0.741 | Open - Forest | -1.77 | 0.174 |

**Table S5.**
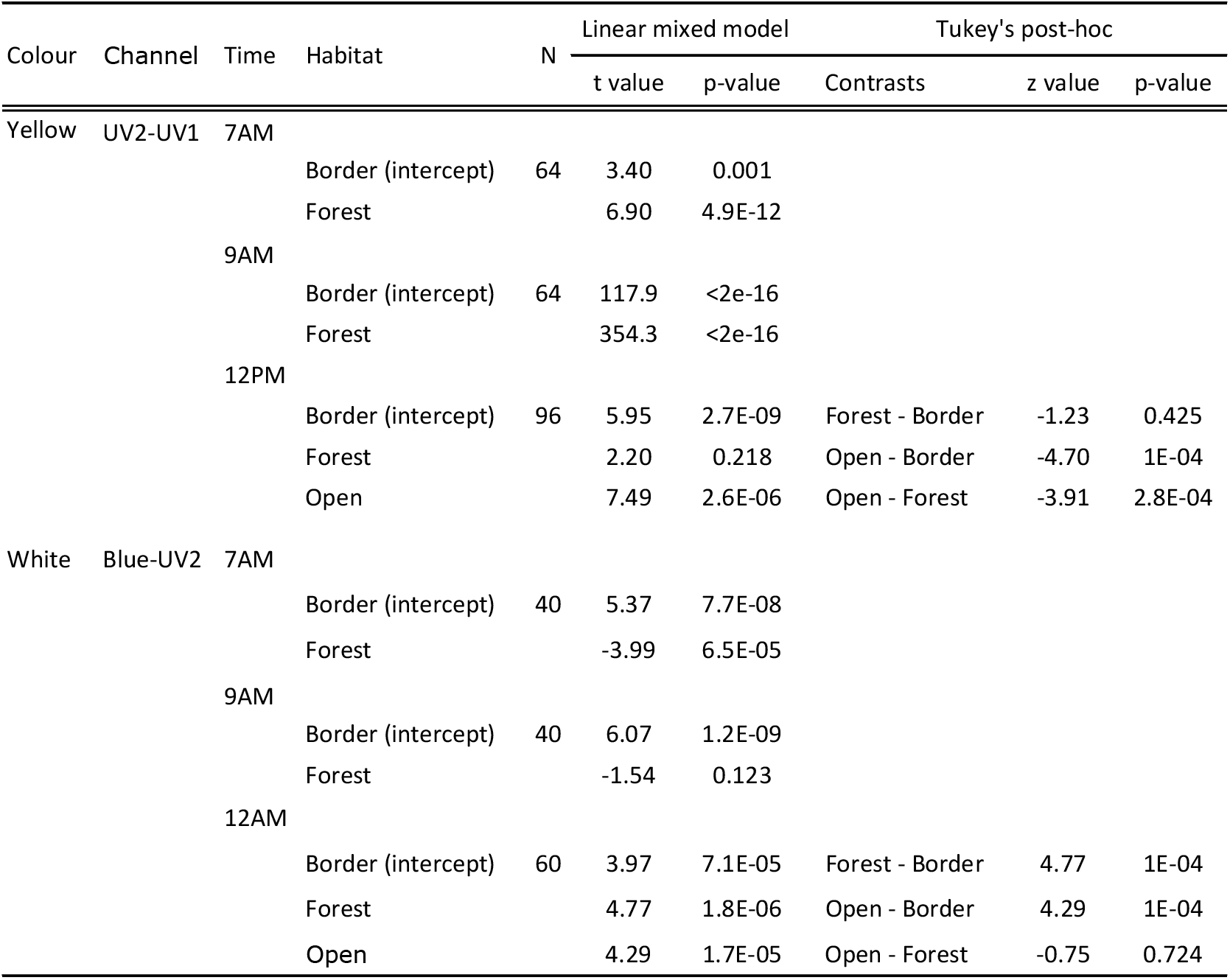
Habitat contrast differences for *Heliconius* vision for yellow and white colours, per opponent channel and time. Generalized linear mixed-effects models with random effects and Tukey’s post-hoc (y ~ Habitat + (1 | Individuals) + (1 | Side)).

**Table S6.** Side of the wing contrast differences for *Heliconius* vision for the red colour, per opponent channel. Generalized linear mixed-effects models with random effects and Tukey’s post-hoc (y ~ Side + (1 | Individuals) + (1 | Habitat:Time)).

| Opponent Channel | Side of the wing | N | Linear mixed model |  |
| --- | --- | --- | --- | --- |
|  |  |  | t value | p-value |
| Red-Green |  |  |  |  |
|  | Dorsal (intercept) | 224 | 6.16 | 7.2E-10 |
|  | Ventral |  | 10.10 | < 2E-16 |
| Blue-Yellow |  |  |  |  |
|  | Dorsal (intercept) | 224 | 6.21 | 5.3E-10 |
|  | Ventral |  | 12.08 | < 2E-16 |
| Blue-UV2 |  |  |  |  |
|  | Dorsal (intercept) | 224 | 15.91 | < 2E-16 |
|  | Ventral |  | 12.90 | < 2E-16 |
| UV2-UV1 |  |  |  |  |
|  | Dorsal (intercept) | 224 | 10.58 | < 2E-16 |
|  | Ventral |  | -1.71 | 0.086 |

